# Behavior of homing endonuclease gene drives targeting genes required for viability or female fertility with multiplexed guide RNAs

**DOI:** 10.1101/289546

**Authors:** Georg Oberhofer, Tobin Ivy, Bruce A. Hay

## Abstract

A gene drive method of particular interest for population suppression utilizes homing endonuclease genes (HEGs), wherein a site-specific nuclease-encoding cassette is copied, in the germline, into a target gene whose loss of function results in loss of viability or fertility in homozygous, but not heterozygous progeny. Earlier work in *Drosophila* and mosquitoes utilized HEGs consisting of Cas9 and a single gRNA that together target a specific gene for cleavage. Homing was observed, but resistant alleles, immune to cleavage, while retaining wildtype gene function, were also created through non-homologous end joining. Such alleles prevent drive and population suppression. Targeting a gene for cleavage at multiple positions has been suggested as a strategy to prevent the appearance of resistant alleles. To test this hypothesis, we generated two suppression HEGs, targeting genes required for embryonic viability or fertility, using a HEG consisting of CRISPR/Cas9 and guide RNAs (gRNAs) designed to cleave each gene at four positions. Rates of target locus cleavage were very high, and multiplexing of gRNAs prevented resistant allele formation. However, germline homing rates were modest, and the HEG cassette was unstable during homing events, resulting in frequent partial copying of HEGs that lacked gRNAs, a dominant marker gene, or Cas9. Finally, in drive experiments the HEGs failed to spread, due to the high fitness load induced in offspring as a result of maternal carry over of Cas9/gRNA complex activity. Alternative design principles are proposed that may mitigate these problems in future gene drive engineering.

**Significance statement:** HEG-based gene drive can bring about population suppression when genes required for viability or fertility are targeted. However, these strategies are vulnerable to failure through mechanisms that create alleles resistant to cleavage, but that retain wildtype gene function. We show that resistance allele creation can be prevented through the use of gRNAs designed to cleave a gene at four target sites. However, homing rates were modest, and the HEGs were unstable during homing. In addition, use of a promoter active in the female germline resulted in levels of HEG carryover that compromised the viability or fertility of HEG-bearing heterozygotes, thereby preventing drive. We propose strategies that can help to overcome these problems in next generation HEG systems.

## Introduction

Gene drive occurs when particular alleles are transmitted to viable, fertile progeny at rates greater than those due to Mendelian transmission. The possibility of using gene drive to bring about population suppression has long been of interest. One such strategy creates conditions in which super-Mendelian transmission results in the spread of a fitness cost through the population. Homing endonuclease genes (HEGs) have been proposed as a vehicle for creating, and at the same time spreading, such a fitness cost (1). A HEG encodes a site-specific endonuclease. When the HEG is located within its target site, in a heterozygous individual, HEG-induced DNA double strand breakage (DSB) at the target site on the wildtype chromosome can lead to repair through homologous recombination, in which the HEG-bearing chromosome is used as the template for repair. This results in the HEG being copied into the broken chromosome in a process referred to as ‘homing’, thereby resulting in an increase in HEG frequency. Naturally occurring HEGs use this transmission distortion mechanism to spread through populations. These HEGs utilize a variety of methods, such as inteins and self-splicing introns, to bring about copying into highly conserved sequences without altering the function of the (invariably essential) genes into which they insert (2). In this way they are able to spread in a population with little or no cost to those who carry them.

Burt proposed that HEGs could also be used to bring about population suppression by spreading a fitness cost through the population (1). In this scenario, homing would be limited to the germline and bring about disruption of genes whose homozygous, but not heterozygous loss-of-function resulted in inviability or sterility. Modeling suggests that such drive elements could spread, and as a result bring about population suppression under a variety of conditions (1, 3–5). However, because homing requires the targeting and cleavage of a specific sequence, its efficacy is sensitive to genomic sequence variation. Variation can occur as pre existing sequence polymorphisms in a population. It can also arise from mutation, and as a result of break repair through non-homologous end joining, which is error prone (6, 7). Regardless of the mechanism, sequence variants that are not cleaved are resistant to homing, and may retain some or complete wildtype gene function. The presence of such resistant alleles can block HEG spread and thereby prevent population suppression (3, 4, 8–11). Thus, the question of how to bring about high frequency homing that is gene specific, but insensitive to some level of sequence variation within the gene, is central to the development of of HEG-based population suppression technologies.

Germline homing and transmission of naturally occurring and engineered HEGs into artificial target sites, and of engineered HEGs into endogenous sites, has been recently demonstrated in mosquitoes and *Drosophila* (7, 12–19). In particular, a number of recent experiments have demonstrated significant rates of germline homing using HEGs created using the CRISPR/Cas9 endonuclease system, in which the Cas9 endonuclease is targeted to specific sequences through association with an independently expressed guide RNA (gRNA). The gRNA includes a ∼20-nucleotide protospacer sequence that mediates RNA/DNA base pairing-dependent target selection. Target sequence limitations with Cas9 are very modest, and thus Cas9 and gRNAs be used to uniquely target most positions in any genome, making them ideal tools for HEG engineering (20–23).

Multiplexing of gRNAs in a Cas9-based HEG so as to target multiple positions within a gene has been suggested as a way to overcome the problem of resistant allele formation (10, 23). The importance of this problem is highlighted by several observations. First, sequence polymorphisms are common in some populations, such as those of mosquitoes (24, 25). Second, germline homing experiments that relied on cleavage at a single Cas9 target site (one gRNA) resulted in the NHEJ-mediated creation of resistant alleles at high rates in the germline, and in zygotes as a result of maternal carryover of active Cas9/gRNA complexes (11, 14–17). The consequences of this were demonstrated in homing based population suppression experiments carried out in mosquitoes, in which genes required for female fertility were targeted by a Cas9-based HEG. HEG frequency initially increased, but this was accompanied by the creation of resistant alleles. Over multiple generations these increased in frequency while those of the HEG decreased (9).

Here we report the development and behavior of two HEGs designed to bring about population suppression by homing into a gene required for embryonic viability or female fertility. In order to test the hypothesis that resistant allele formation can be prevented by gRNA multiplexing each HEG included four gRNAs designed to cleave the gene of interest. We determined the homing and cleavage rates of these elements, and characterized the products generated post cleavage on a molecular level. Cleavage rates were high. Homing rates were in general modest, varied greatly between individuals and were associated with HEG instability. Importantly however, cleavage resistant alleles were not observed. Finally, in drive experiments we observed that after an initial increase in allele frequency, the HEGs proceeded to drop out of the populations. Possible reasons for this are discussed.

## Results

### HEG design

We selected two target genes for our HEG constructs, *yellow-g* and *deformed* (*Dfd*). The *yellow-g* gene is somatically expressed in follicle cells of the ovary and encodes a protein required for egg shell formation. Loss of *yellow-g* results in female sterility, due to collapse of laid eggs (26). The *Dfd* gene encodes a HOX family transcription factor expressed in the early embryo head anlagen, and is essential for posterior head morphogenesis, with loss of *Dfd* resulting in embryonic lethality (27). Each HEG included Cas9 and four gRNAs (Fig. 1A). Cas9 expression was driven by a *nanos* regulatory element, which drives germline expression in males and females (28). This transgene was flanked by gypsy insulators to ensure germ line specific expression. The gRNA expression constructs consisted of two tandem pairs, in which one gRNA of each pair was expressed under the control of the *Drosophila* U6:3 regulatory sequences, and the other was expressed under the control of U6:1 regulatory sequences (29). To increase gRNA expression levels, the gRNA scaffolds were modified as detailed previously, to eliminate cryptic termination sequences (30). One U6:3-gRNA/U6:1gRNA tandem was placed to the left of Cas9, while the other was placed to the right. Homology arms of approximately 1kb were included to the left and right of the two outermost gRNAs to promote homologous recombination. Finally, the HEG contained a *3xp3-td-tomato* dominant marker (Fig. 1A).

**Fig. 1:**
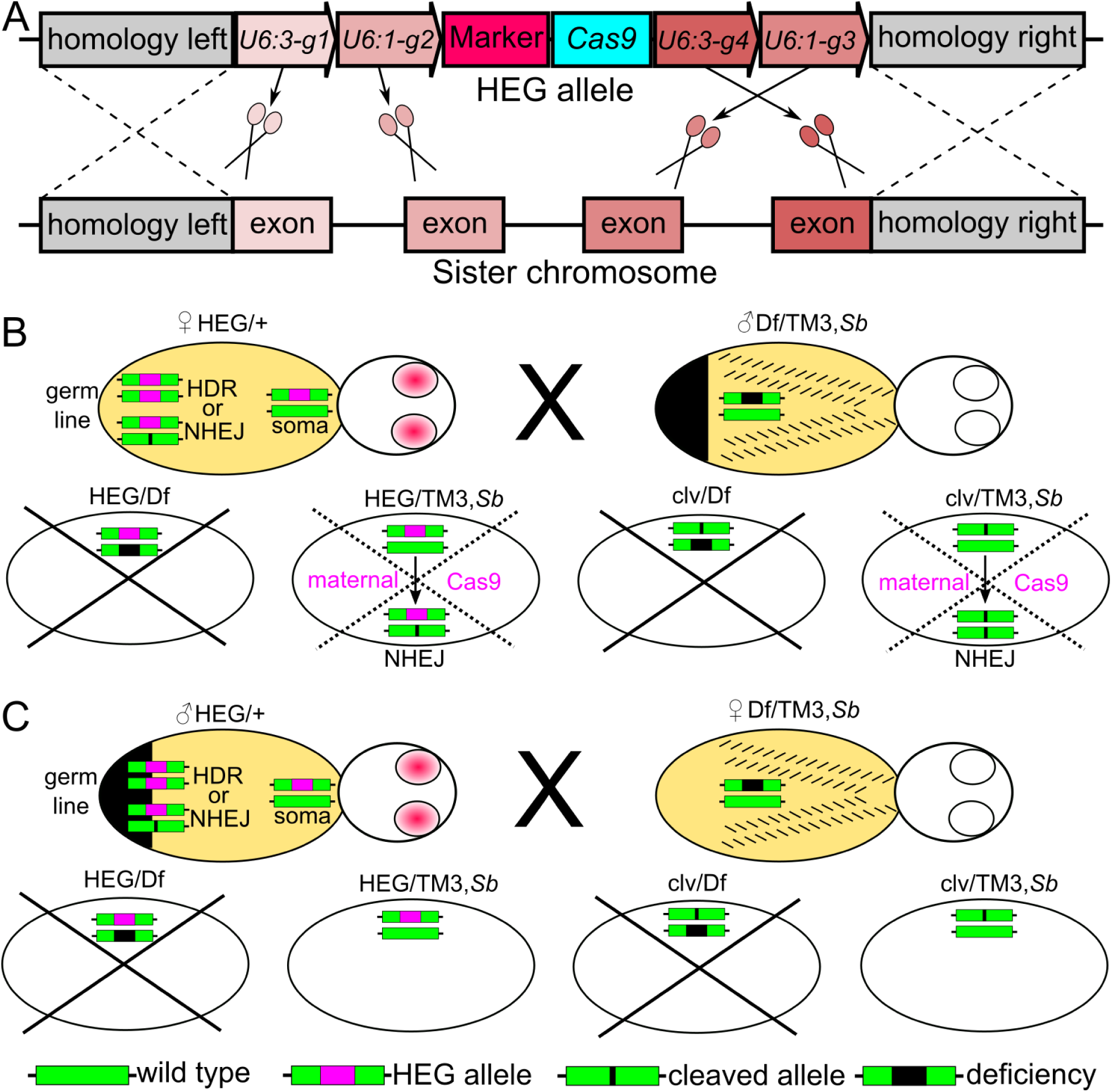
**HEG design and possible cross outcomes for the HEG targeting *Dfd* (A) Design of CRISPR based HEG construct.** The HEG element consists of two outer homology arms to mediate HDR with the wild type allele on the homologous chromosome. Between the arms are a set of four U6 (alternating U6:3 and U6:1) driven guide RNAs, Cas9 driven by germ line specific *nanos* regulatory elements, and a *3xp3-td-tomato* marker. **(B-C) Cross outcomes for HEG heterozygote for the *Dfd*-HEG crossed to a deficiency line, with the HEG coming from a female (B) or male (C)**. Shown are parents of the crosses (upper row) and genotypes of progeny (ovals). In the germ line the wild type allele was either converted to the HEG allele by HDR or to a cleaved allele (clv) via NHEJ or incomplete homing. Parental genotypes were HEG heterozygotes (HEG/+) and heterozygotes for a stock carrying a deficiency that removes *Dfd* or *yellow-g* and the third chromosome balancer TM3 (Df/TM3, *Sb*). Genotypes of the offspring were HEG allele over deficiency (HEG/Df), HEG allele over balancer (HEG/TM3, *Sb*), cleaved allele over deficiency (clv/Df), cleaved allele over balancer (clv/TM3, *Sb*), non cleaved allele over deficiency (+/Df, not shown), and non cleaved allele over balancer (+/TM3, *Sb*; not shown). Genotypes of interest in the embryos are depicted as ovals, lethal genotypes (or sterile female genotypes in the case of *yellow-g*) are crossed out (dashed crosses if due to maternal carry over). (B) When the HEG was transmitted through the female germline, all embryos had Cas9 and gRNA deposited during oogenesis, leaving all embryos with active HEG components. (C) No paternal carry over of Cas9 to the embryos was observed: all the offspring that inherited a wild type allele from the deficiency strain (from the balancer TM3, *Sb*) were viable in the case of the *Dfd*-HEG, or fertile with the *yellow-g*-HEG (*yg*-HEG).

### Homing and cleavage rates of HEG strains

Homing, and cleavage without homing, result in the creation of loss-of-function alleles of the target genes. We scored for the presence of these events by crossing HEG heterozygous males and females (HEG/+) to flies heterozygous for a deficiency (Df) that removes the target locus and a version of the TM3, *Sb* balancer chromosome carrying the dominant marker *Sb* (Df/TM3, *Sb*). By comparing the frequency of specific outcomes in progeny that carried the Df chromosome versus the TM3, *Sb* balancer chromosome we were able to distinguish between homing and cleavage events. The presence of homing events, or the original HEG chromosome, was inferred by the presence of the *td-tomato* dominant marker. Cleavage without homing was inferred based on the presence of a loss-of-function phenotype in the absence of the *td-tomato* marker (see Fig. 1B,C). To get an overview of the frequency of each event type and inter-individual variability, we set up 25 single fly crosses of each sex for both of the HEG constructs. Homing rates were calculated from the relative proportion of HEG/TM3, *Sb* to (+ or clv)/TM3, *Sb* progeny. Cleavage rates were calculated based on the frequency of surviving or fertile +/Df individuals, which represent either resistant alleles or non-cleaved alleles. As discussed below, molecular analysis showed that no completely resistant (lacking target sites for all four gRNAs) alleles were formed (see Materials and Methods for details and formulas and Table S1 for counts).

When the *Dfd*-HEG came from a mother, most progeny died, presumably due to maternal carry over and subsequent cleavage of *Dfd.* With this low number of surviving progeny, it was not possible to calculate meaningful homing rates (see Table S1 for counts). For the remaining crosses, results are shown in Fig. 2A-C. Interestingly, homing rates for both HEGs showed high variability between individual single fly crosses, ranging from 0 to 83%, while cleavage rates were consistently very high (average homing and cleavage rates as mean (standard deviation): ♂*Dfd*-HEG/+ homing= 0.33(0.21), cleavage= 0.99(0.02); ♂*yg*-HEG/+ homing= 0.19(0.17), cleavage= 1(0.00); ♀*yg*-HEG/+ homing= 0.26(0.20), cleavage= 0.89(0.31)).

**Fig. 2:**
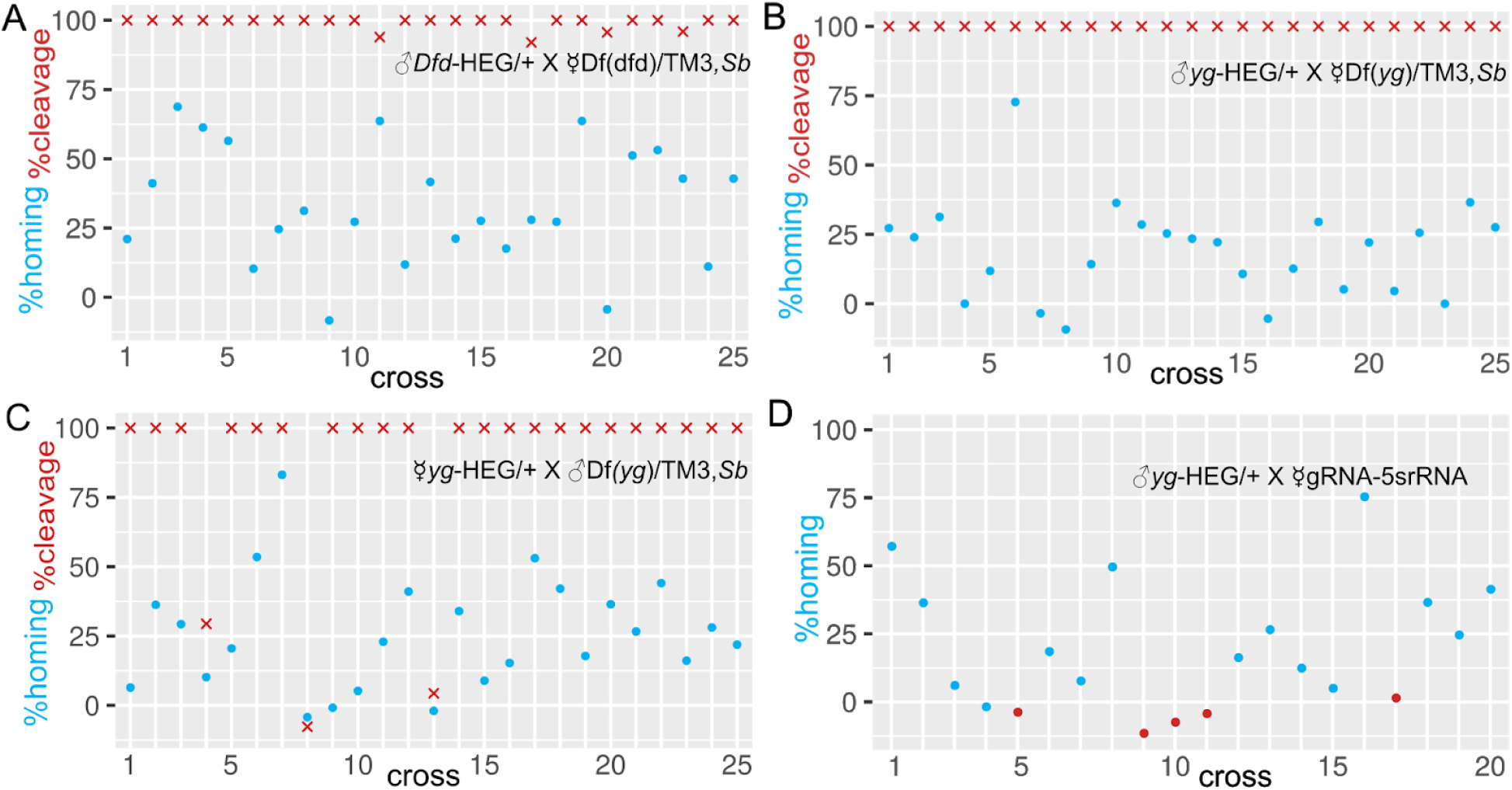
**(A-C) Homing- and cleavage rates of HEG bearing heterozygotes crossed to a deficiency line.** Homing rates are denoted as blue dots, cleavage rates as red crosses on the y-axis, with the number of each individual cross indicated on the x-axis. Homing rates were highly variable between crosses, whereas cleavage rates were 100% for most of the crosses. **(D) Test crosses to determine Cas9 activity in the *yg*-HEG.** Shown are the homing rates of heterozygous *yg*-HEG males crossed to a gRNA strain targeting the 5s rRNA that results in sterility in all offspring when crossed to a strain with functional germline specific Cas9. Homing rates varied similar to the crosses to the deficiency line. In 5 out of 20 crosses the progeny were fertile (red dots) and no homing occurred, indicating that Cas9 in the HEG strain had lost functionality, even though the HEG cassette dominant *td-tomato* marker was still present (Note that since homing is calculated based on deviation of progeny genotype ratios from those of Mendelian transmission, homing rates of less than zero can occur. These likely reflect cases in which no homing occurred, and genotypes due to Mendelian segregation were not balanced.

### Fitness costs from germline carry over

In order for suppression HEGs to spread, cleavage and homing must be restricted to the adult germline. Otherwise heterozygotes and/or their progeny will show loss-of-function phenotypes, which by definition include a fitness cost. Maternal carryover of Cas9/gRNA complexes into non-Cas9-bearing individuals has been shown to result in high frequency cleavage of target sequences in *Drosophila* (17, 31). Similar effects can also be inferred from the results of other work in *Drosophila* and mosquitoes that utilized promoters driving Cas9 in the female germline (14–17). The frequency of cleavage in the zygote due to expression in the male germline is less clear, but likely to be low (32). Here we address the significance of carryover in the context of HEGs designed to induce a fitness cost.

For the *Dfd*-HEG, we were unable to determine the extent of male carryover-dependent cleavage in the zygote because progeny genotypes that include a homed or cleaved paternal allele and a cleaved maternal allele die during embryogenesis. For the *yg*-HEG we determined the frequency of carryover-dependent cleavage from the germline of male *yg*-HEG/+ individuals crossed to Df/TM3, *Sb* females by determining the frequency of TM3, *Sb* bearing female progeny (*yg*-HEG/TM3, *Sb*; +/TM3, *Sb* and clv/TM3, *Sb*) that were sterile. None should be sterile if carryover-dependent cleavage did not occur. 141 TM3, *Sb* female progeny of the first 5 crosses of Fig. 2B were tested. Of these, only 3 did not produce offspring (2.1%), suggesting that carryover-dependent cleavage is minimal. Based on this observation we assume below that carryover-dependent cleavage in the zygote is female-specific.

For the *Dfd*-HEG we estimated fitness cost indirectly, under the assumption that crosses between HEG/+ and Df(*Dfd*)/TM3, *Sb* adults would, in the absence of carryover into the embryo, produce equal numbers of progeny regardless of which parent carries the HEG or the Df for the locus. With this assumption, and in the absence of carryover, the number of TM3, *Sb*-bearing progeny (HEG/TM3, *Sb* & +/TM3, *Sb*) should be similar, regardless of whether the HEG came from the mother or father. In actuality, when the HEG came from the mother the total number of adult TM3, *Sb*-bearing progeny from all 25 crosses of Fig. 2 was 97 and large numbers of dead eggs were observed. In contrast, when the HEG came from the male 1218 TM3, *Sb* progeny were obtained, suggesting a maternal carryover-dependent fitness cost of 92%. In the case of the *yg*-HEG, we calculated the fitness cost due to maternal carry over directly, as the number of sterile females (HEG/TM3, *Sb* and clv/TM3, *Sb*; 405) divided by the sum of all females with the balancer chromosome (HEG/TM3, *Sb* & clv/TM3, *Sb* & +/TM3, *Sb*; 489), suggesting a maternal carryover-dependent fitness cost of 82.8%. Together, these results support earlier observations (17) and demonstrate that when using the *nanos* regulatory sequences maternal carryover-dependent cleavage of paternal alleles in the zygote is common, and that fitness costs to HEG-bearing zygotes is high. The consequences of these fitness costs for gene drive are explored further below.

### Molecular characterization of cleavage events

Because most progeny of *Dfd*-HEG-bearing heterozygous mothers died during embryogenesis as a result of carryover-dependent cleavage, we were unable to further characterize these individuals. However, since *yellow-g* is only required for eggshell formation in somatic ovarian cells of adult females, female clv/Df(*yg*) offspring of the ☿*yg*-HEG/+ X ♂Df(*yg*)/TM3, *Sb* cross and the ♂*yg*-HEG/+ X ☿Df(*yg*)/TM3, *Sb* cross are viable, but sterile. Genomic DNA was isolated from 10 sterile female cross progeny (two flies each from five different vials) where the HEG came from a female (F1-10), and 10 where it came from a male (M1-10). All flies analyzed lacked the *td-tomato* marker, indicating that a complete homing event had not occurred. However, these females were sterile, suggesting their genotype was clv/Df. The target region was amplified by PCR and sequenced. Since the Df-bearing chromosome lacks the *yellow-g* locus, all products must come from a cleaved or partially homed locus.

We found a wide range of different cleavage events (Fig. 3). Nine flies showed deletions between the four gRNAs. In this category, five flies had a deletion between the outer gRNA1 and gRNA4 (F1, F3, F8, M5 with additional bases deleted, and M7). One had a deletion between gRNA1 and gRNA3 (M1 with PAM of gRNA4 mutated). Two had a deletion between the intron located between gRNAs 1 and gRNA2 and gRNA4 (M3, M4, coming from the same parents), and one had a deletion between gRNA2 and gRNA3 (M6). Finally, one fly had only a small deletion at the gRNA1 target site (M2, also had a mutated PAM for gRNA4).

**Fig. 3:**
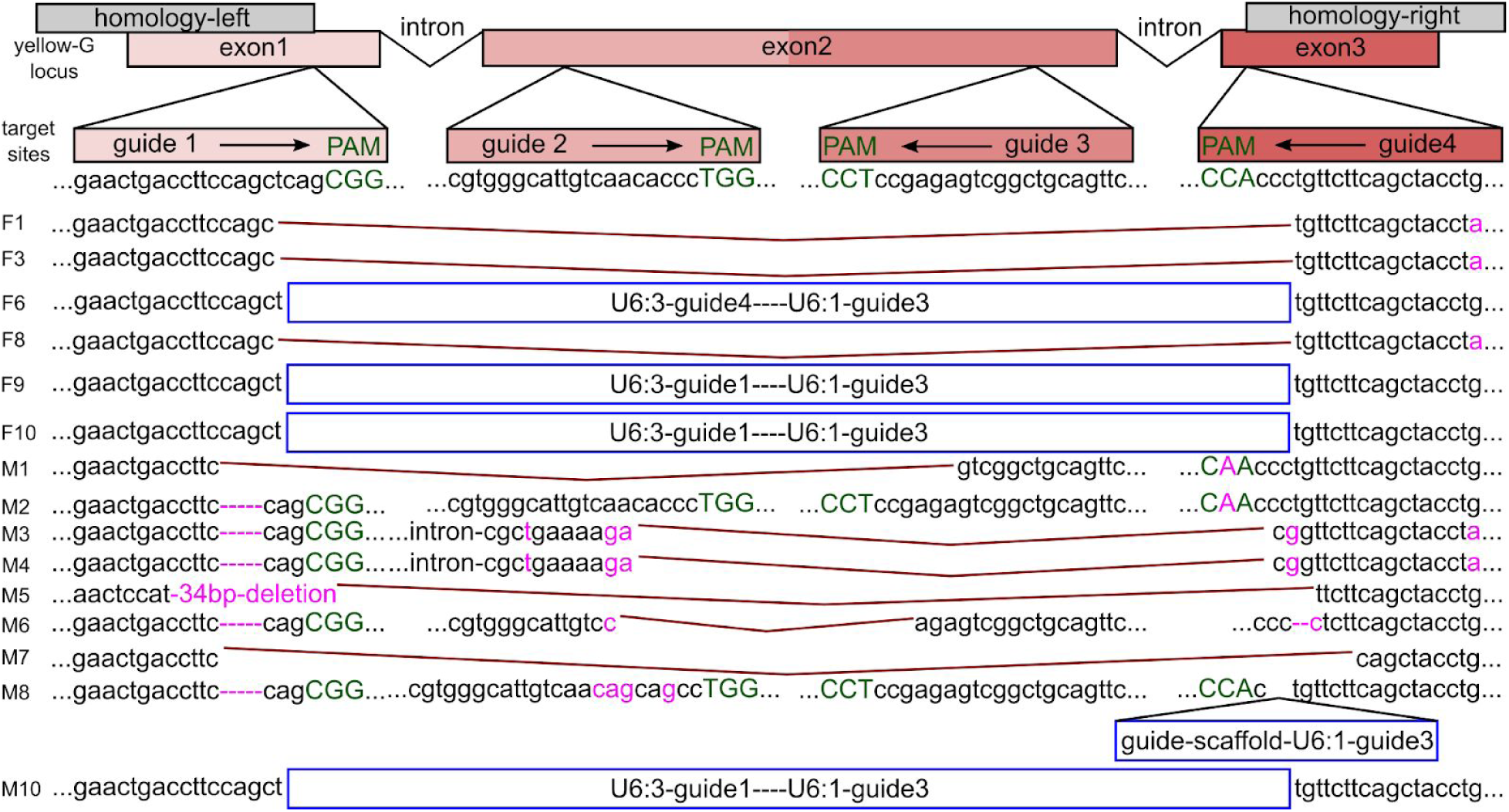
**Cleavage events.** Sanger sequencing results from individual sterile progeny of the clv/Df(*yg*) genotype, in which progeny come from ☿*yg*-HEG/+ x ♂Df(*yg*)/TM3, *Sb* (F1-F10) or from ♂*yg*-HEG/+ x ☿Df(*yg*)/TM3, *Sb* (M1-M10). Flies were taken in pairs from individual crosses, i.e. F1 and F2 coming from one vial, F3 and F4 from another and so on. PAM in green, mutations and small deletions in magenta, large deletions between gRNA target sites are depicted with brown lines and HEG cassette remnants due to incomplete homing in blue.

For the remaining ten flies analyzed, we detected incomplete or partial homing events. Three of these had half of the left and half of the right guide cassette inserted between gRNA1 and gRNA4 cut sites (F9, F10, and M10). One had the complete right guide cassette inserted between gRNA 1 and gRNA4 (F6). One had half of the right guide cassette inserted into the cut site of gRNA4 (M8). For the remaining five flies we were not able to generate PCR products that spanned the two outer cut sites. However, we could amplify a part of the Cas9 sequence. Since these flies lacked the *td-tomato* marker, but retained other parts of the HEG cassette (at least some of Cas9), they were also considered to represent incomplete homing events (F2, F4, F5, F7, and M9, not shown). Together these results show that the consequences of Cas9-mediated cleavage are diverse, and include a significant frequency of incomplete homing events.

### HEG cassette stability

Given the high frequency of incomplete homing events, wherein the *td-tomato* marker was lost but Cas9 was still present, we were interested to determine if incomplete homing events also resulted in events in which Cas9 was lost, but the *td-tomato* marker was retained. To test this possibility we isolated males that were *td-tomato* positive (*yg*-HEG-bearing) at generation 10 from a drive experiment (below), in which the *yg*-HEG had been introduced into a wildtype population at a frequency of 25%. These males were outcrossed to *w*-females to ensure heterozygosity for the HEG. Twenty male heterozygous progeny from these crosses were mated to females of a Cas9 tester strain that expresses under the control of U6:3 regulatory sequences a gRNA designed to bring about cleavage of the 5s ribosomal RNA repeats (gRNA-5srRNA) (Fig. 2D). When gRNA-5s females are crossed to males expressing Cas9 under the control of *nanos* regulatory sequences, all offspring are sterile, showing that sterility can be used as a test for the presence of functional Cas9. Progeny from the above 20 crosses were scored for the *td-tomato* marker to determine homing rates, and all females were outcrossed to *w*-males to score for fertility. The results are shown in Fig. 2D (counts in Supplementary Table 1). Female progeny of 5 of the 20 crosses were fertile indicating that Cas9 function had been lost. As expected, no homing was observed in these crosses. The rates of homing for the remaining crosses varied within a range similar to the one observed in crosses to the deficiency-bearing lines (Fig. 2A-C).

### Resistant alleles are absent among escapers

As discussed above and in Fig. 2, the cleavage rates of target loci in the male and female germline of HEG/+ adults was very high for both *Dfd*-HEG and *yg*-HEG. However, in rare cases the cross between HEG/+ and Df/TM3, *Sb* adults resulted in progeny that carried the Df chromosome (+/Df or clv/Df), but were viable (*Dfd*) or fertile (*yellow-g*). To determine whether these individuals simply carry alleles that escaped cleavage, or alleles that are resistant to cleavage, we isolated genomic DNA and carried out PCR and sequencing of the target region for all of them (Table 1 and Fig. 4). For the *yg*-HEG, 24 female escapers (fertile) were isolated from three crosses in which the HEG came from the female (F4, F13, and F8). The six escapers from cross F4 had a mutated PAM at the gRNA4 target site; the 11 escapers from cross F13 had a mutation at base 1 of the gRNA4 target site; the seven escapers from cross F8 had an intact target gRNA4 target site. All other target sites were intact in all escapers.

**Fig. 4:**
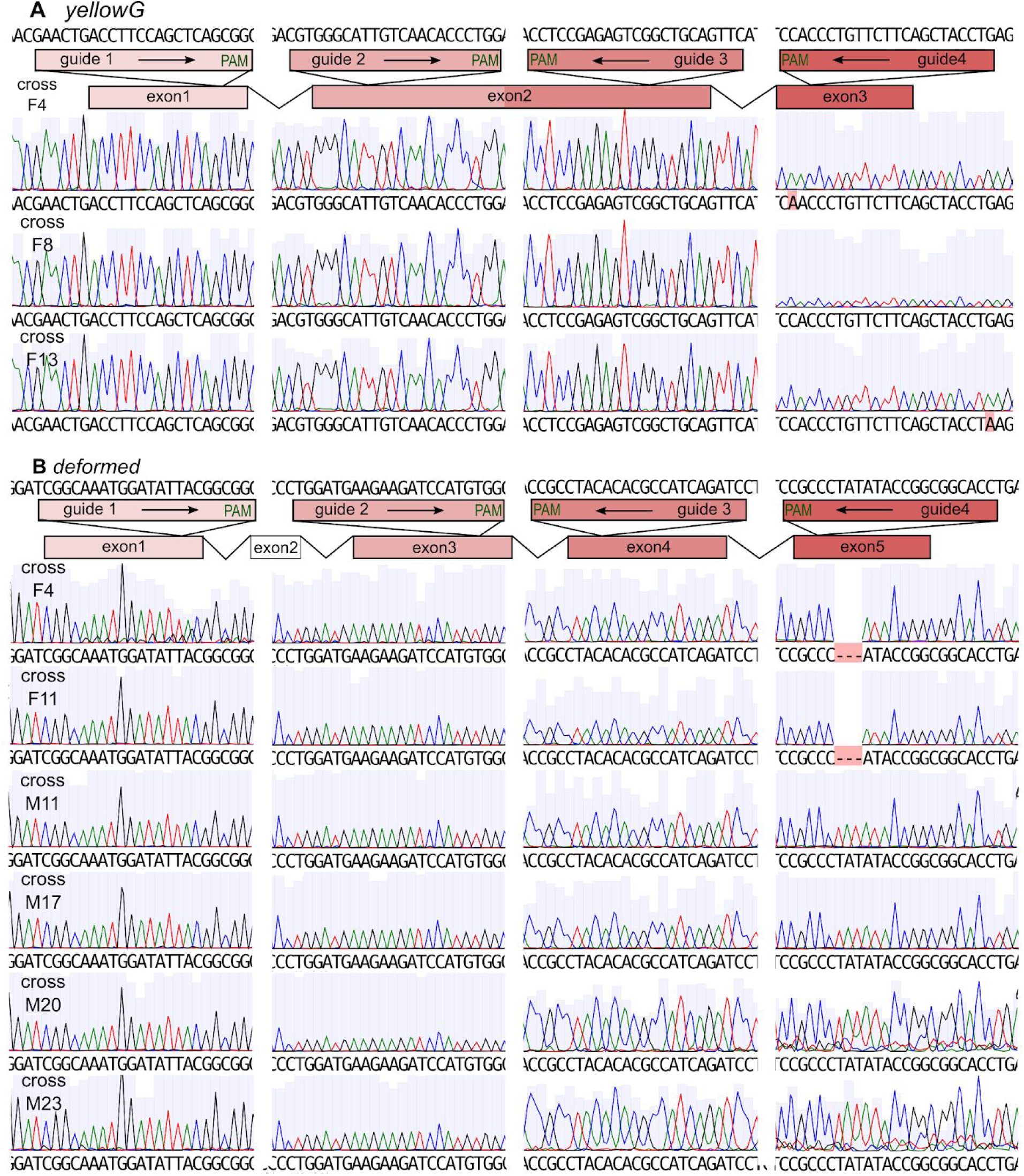
**gRNA target site sequences of escapers.** Chromatograms from one fly of each cross containing escapers is shown. All flies were of the +/Df genotype **(A) *yg*-HEG Escapers.** gRNA4 target sequence had a mutated PAM site in progeny of cross F4. Progeny from cross F13 had a mutation in the distal most base of the PAM. All other target sites did not show mutations. **(B) *Dfd*-HEG Escapers.** Progeny coming from cross F4 and F11 had a 3bp deletion in the target sequence. All other target sites were intact.

**Table 1:**
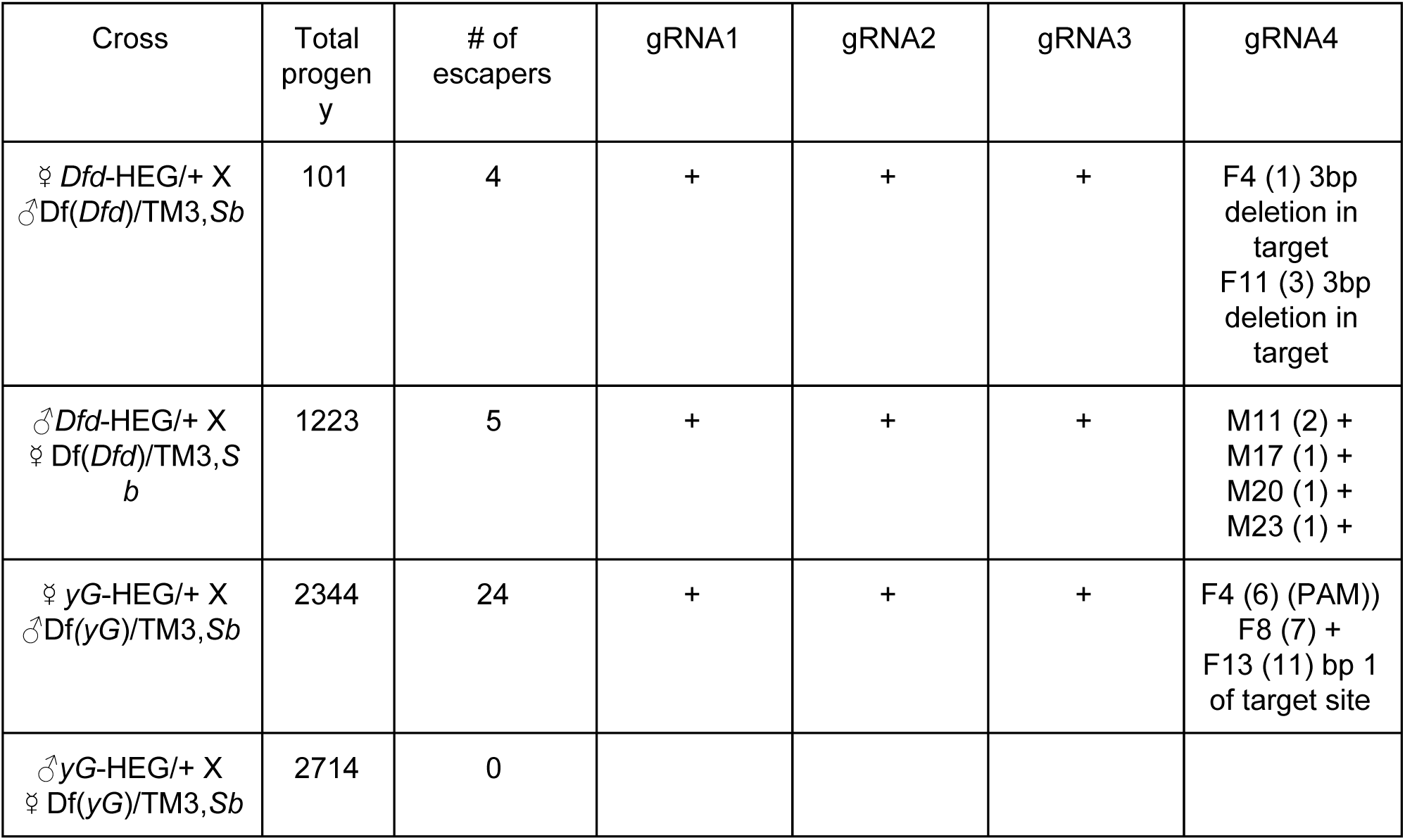
Escaper and resistant gRNA target sites. Shown are the total number of progeny and number of escapers from all 100 single fly crosses to the deficiency lines. Intact gRNA target sites are indicated with “+”. When a mutation was observed, the cross and the number of individuals is indicated (), and a short description of the mutation is provided.

For the *Dfd*-HEG we found four escapers from two crosses where the HEG came from a female (F4 and F11), and in five individuals from four crosses where it came from a male (M11, M17, M20, and M23). Escapers from crosses F4 and F11 had a 3bp in-frame deletion 3bp upstream of the PAM of the gRNA4 target site. All other target sites in the remaining escapers were intact. Together these results make two important points. First, multiplexing prevents the appearance of alleles resistant to cleavage at all four sites. Second, while the use of multiple gRNAs results in a very high frequency of cleavage, it does not completely prevent evasion of cleavage at intact sites. Possible reasons for this are discussed below.

### Gene drive outcomes

To assess the potential of our HEG constructs to suppress populations, we set up several drive experiments. As the seed for generation 0, we introduced *w*-females that had been mated to HEG/+ males, and *w*-females mated to *w*-males. These two groups of mated females were mixed at a 1:1 ratio, corresponding to a HEG-allele introduction frequency of 25%. Four replicates were carried out for each drive experiment. In each generation adults were allowed to lay eggs for two days and then removed. After progeny had matured to adulthood we scored for the presence of the *td-tomato* marker, seeded the next round with 200 flies and repeated the cycle. As a gene drive control we used inactive HEG constructs that were inserted at the target locus and expressed the dominant marker and gRNAs, but that lacked Cas9.

The dynamics of HEG behavior are illustrated in Fig. 5. In the case of *Dfd*, the HEG increased in frequency for one generation, but decreased continuously thereafter. By generation 9 the *Dfd*-HEG had been completely lost from all replicates. In contrast, the inactive*-Dfd*-HEG remained in the population. In the case of *yellow-g*, the HEG increased in frequency for two generations. It then decreased in frequency, as with the *Dfd*-HEG, albeit at a slower rate. By generation 12 the HEG had been lost completely from one replicate, and had fallen below the introgression frequency in the others. The control version of the *yg*-HEG increased in frequency slightly above the introgression frequency up until generation 7, before returning to around 25% at generation 12.

**Fig. 5:**
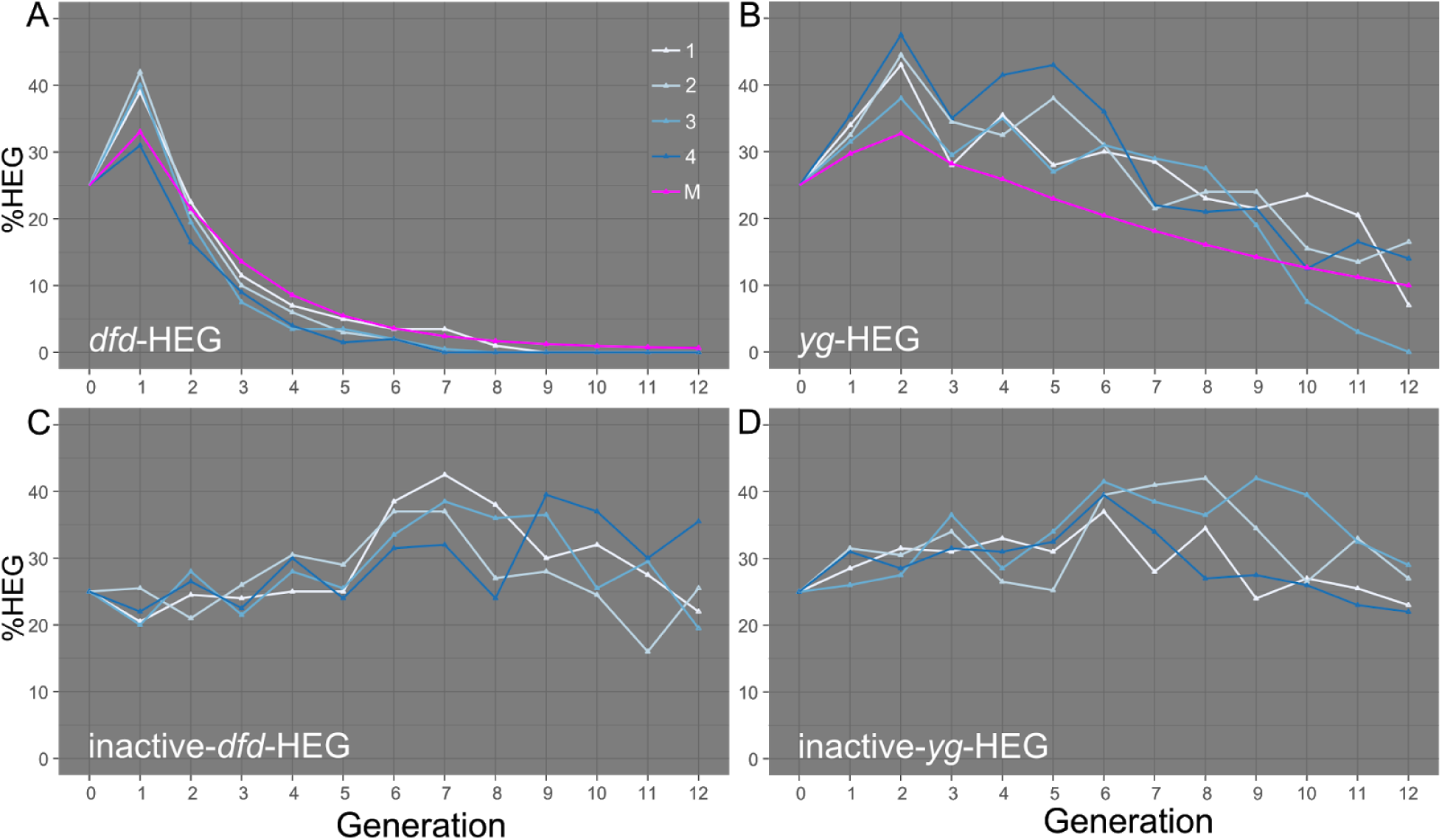
**Drive experiments. (A-D)** Plots of drive experiments The frequency of HEG-bearing individuals (HEG/+ and HEG/HEG) is indicated on the y-axis and the generation number on the x-axis. Drive replicates are labeled as 1-4. Predicted drive behavior, using fitness costs inferred from data is in pink (M).

We generated a deterministic, population frequency model of our two HEGs, incorporating the observed cleavage and homing rates, and fitness costs due to maternal carryover-dependent cleavage (see methods for details). Models for both HEGs fit the data well. In the case of *Dfd*, the model predicted an increase in frequency for one generation followed by a rapid decrease, as was observed (Fig. 5A). In the case of *yellow-g* the model predicted a two-generation increase in the frequency of HEG-bearing individuals before a slow decline, as was also observed in the drive experiments (Fig. 5B). These observations suggest that the variables studied and quantified in this work, with respect to homing, cleavage and carryover, are sufficient to capture major aspects of HEG behavior.

## Discussion

Homing-based gene drive requires the targeting of specific sequences for cleavage, and is therefore sensitive to existing sequence variation or novel sequences created by error-prone repair pathways. Homing based gene drive using Cas9 and a single gRNA has been demonstrated in *Drosophila* (14, 17), mosquitoes (15, 16), and yeast (33). However, these studies also reported the creation of mutant alleles that are resistant to cleavage. In those experiments in which coding regions were targeted for cleavage, resistant alleles included variants in which target gene function was retained (9, 14, 17). The presence of such alleles is predicted to prevent population suppression when genes required for viability or fertility are targeted (3, 10, 11). Resistant alleles can also prevent homing-based population replacement using HEGs that carry a gene of interest into the population for the same reason. Thus, prevention of resistant allele formation is a central problem that must be solved in order for homing to be useful as a drive mechanism.

Multiplexing of gRNAs has been proposed as a way of overcoming this problem, but has not been tested directly (10, 23, 34, 35). In this study we tested the effects of gRNA multiplexing on resistance allele formation in the context of suppression HEGs developed with components used used in earlier gene drive experiments in insects. A key positive finding of our work is that when four gRNAs are used to target a gene, cleavage rates in the germline are very high, and resistance allele formation is prevented. However, our work also highlights several important problems that remain to be solved. First, homing rates in the male and female germline are overall modest, and vary greatly between individuals. Second, HEGs are unstable, with homing resulting in the creation of a high frequency of chromosomes that bear incomplete HEGs, lacking one or more essential elements. Third, maternal carryover of a Cas9/gRNA HEG results in extensive cleavage of the paternal allele in zygotes, and this results in the creation of HEG heterozygotes (clv/HEG) that experience dramatic fitness costs. Together, these results predict that the two HEG generated, *Dfd*-HEG and *yg*-HEG, should fail to spread; drive experiments confirmed these predictions, with both elements initially increasing in frequency, but then decreasing in frequency and/or being lost from the population. Below we discuss these results in more detail and explore possible paths forward.

### Cleavage and homing rates

For the *yg*-HEG, cleavage rates were 100% in the germline of all but three of 50 individuals (Fig. 2B, C). In two of the three fertile females (F8 and F13), cleavage rates were likely zero (homing rates were zero), while in the third (F4), a cleavage rate of ∼26% was inferred. Low rates of cleavage in these individuals could reflect variation in the levels of Cas9, the timing of its expression, expression of gRNAs, or accessibility of the target sites. Another possible reason for low (or no) cleavage in these individuals is suggested by our molecular analysis of homing events, which uncovered a high frequency of incomplete homing. In particular, when we examined flies in which homing of the *yg*-HEG had been permitted over 10 generations in a drive experiment (Fig. 5), 25% were found to have lost Cas9 activity, but retained the dominant marker used to score for the presence of the HEG (see Fig. 4D). Thus, some ostensibly HEG-bearing individuals tested for cleavage, such as F8, in which all target sites are wildtype in sequence, may lack a functional HEG. Multiple female progeny of F4 carried the same mutation in gRNA4; multiple female progeny of F13 also carried a distinct, common mutation in gRNA4. Both mutations were also observed in our analysis of cleavage events in sterile females (Fig. 3). These results suggest that these sequence variants represent pre-existing polymorphisms. Regardless of their origin, the fact that gRNA targets 1-3 were intact in these individuals indicates that cleavage did not occur at all sites. gRNAs 1-3 are likely to be functional, given the spectrum of cleavage products observed in sterile clv/Df females (Fig. 3). It is possible that interaction of gRNAs with target sequences is a relatively slow, stochastic process, with these simply representing the rare alleles in which gRNAs 1-3 did not bind to DNA. It is also possible that gRNAs 1-3 are expressed/processed/loaded and/or promote cleavage less efficiently than gRNA4, which may be important if Cas9 levels are limiting. A possibility of particular concern, consistent with our data, is that gRNA 4 is preferentially loaded. If there are also polymorphisms in the gRNA 4 target site (pre-existing or created through NHEJ) that prevent cleavage, the combination of these two events would result in an abundance of Cas9/gRNA complexes that are effectively dead with respect to the target gene, even though gRNA target sites are intact and the gRNAs expressed. Our data on escapers of cleavage by the *Dfd*-HEG, while representing small numbers, may be more suggestive of a situation in which multiple gRNAs are often utilized. Thus, in contrast to the case of *yellow-g*, in which cleavage rates were low in females that gave rise to escapers (F4, F8, F13), the four *Dfd*-HEG/+ males that gave rise to +/Df viable progeny still had very high cleavage rates (only one or two viable progeny/adult). The five rare escapers progeny from these four males were wildtype in sequence at all target sites, consistent with models in which cleavage simply failed to occur at any of the four sites some of the time. Little is known about how specific gRNAs are chosen from a cellular pool, and what regulates the kinetics of subsequent steps. Much remains to be learned about how to ensure contemporaneous if not simultaneous loading and cleavage at multiple sites within a gene. The hypothetical scenario outlined above involving preferential loading of a gRNA targeting a sequence that can mutate to resistance demonstrates how easily even multiplexing based strategies to prevent resistance development could be defeated. Exploration of these and other variables such as the levels of gRNAs and Cas9, nuclear import of Cas9, stability of particular gRNAs, and accessibility of specific loci in the germline should provide guidance in how maximize opportunities for multiple cleavage events.

In contrast to the generally very high rates of germline cleavage of target loci, homing rates for individual *yg*-HEG males and females, and *Dfd*-HEG males, showed great variability, varying from 0-83% (mean values: ♂*Dfd*-HEG=0.33; ♂*yg*-HEG=0.19; ♀*yg*-HEG=0.26). Some variability in homing rate between individuals has also been noted previously (17) in experiments targeting the *y* locus, though the variability we observe is particularly high. The average homing rate we observed is also lower than that observed by Champer (17). The basis for this variability is unknown, but important to understand as it seems generally unrelated to the frequency of cleavage, which is required for homing to occur. Very high homing rates were inferred in one earlier *Drosophila* study in which *y* was targeted with a HEG that lacked a dominant marker with which to follow homing independently of mutation of the *y* locus (14). However, the vasa promoter used in these experiments results in extensive maternal carryover, and is active in somatic cells (17), leaving open the possibility that the high rates of homing inferred may reflect, to some extent, cleavage in somatic cells of heterozygotes rather than homing.

One possible explanation for the high variability we observe in homing is that having multiple Cas9/gRNA complexes bound to to the locus (but before cleavage) interferes with some aspect of homologous recombination-dependent repair initiated following cleavage at one of these sites. For example, it has been suggested that binding of one Cas9/gRNA complexe could interfere with binding to a neighboring site as a result of DNA supercoiling (36). Such a mechanism might contribute the great variability in homing rates, since which combination of target sites is occupied at any one time in a specific germ cell, and what the order of cleavage is, will vary. If such an effect does exist, it may be possible to overcome it by increasing the distance between gRNA targets and/or changing their orientation (though other possible effects, discussed below, may argue for placing cleavage sites near each other). The timing of Cas9 and gRNA expression may also matter. While we observe very high frequencies of cleavage, which indicates that Cas9 and gRNAs are expressed and active, the exact timing of this expression and its variability are unknown. The time at which cleavage occurs is likely to be important for the choice of repair pathway. It may also be important that target loci for suppression HEGs such as *Dfd* and *yellow-g* are by definition not important for germline development, since otherwise homing (unless it occurred after gene function was required), would result in loss of cells in which homing occurred. *Dfd* and *yellow-g* are not expressed at appreciable levels in the germline. Thus, it is also possible that lack of expression creates a chromatin environment in which homing steps subsequent to cleavage are rendered challenging. Finally, we note that in a multiplex design, the sequences on either side of a cut site often do not correspond to those present in the homology arms flanking the HEG (Fig. 1). They do in single gRNA designs (14–17); they also do when both outer gRNAs cleave in our multiplex design (Fig. 1); however, only one (gRNA1 alone or + gRNA2/gRNA3; gRNA4 alone or + gRNA2/gRNA3) or no ends (gRNA2 and/or gRNA3) correspond to the homology arms in other cleavage scenarios. Our gRNAs span a region of 2.2kb within the *yellow-g* gene and 8.8kb within the *Dfd* locus. Thus, ends generated by cleavage at any combination of gRNAs other than the combination of gRNA1 and gRNA4 presumably require extensive resection before HR-dependent repair can occur. This may reduce the frequency and/or efficiency of HR versus other competing repair pathways in copying a complete HEG. If the location of the break with respect to the location of homology arms is critical, it will be important to understand how to locate sites - perhaps close together - so as to minimize negative consequences for HR-based repair.

### HEG instability for a suppression HEG

In our crosses to the deficiency lines we observed the homing rate to be highly variable between replicates, and low on average (19-33%). When we assayed flies carrying the *td-tomato* marker, we found that 5 out of 20 tested had lost Cas9 activity after ten generations in a drive experiment (Fig. 2D). One important implication of this observation is that scoring for the presence of a dominant marker located within the HEG provides only an upper estimate on the rate of complete homing events. To mitigate this problem it may be possible to link Cas9 and the dominant marker into a single dicistronic transcript by making use of 2A-like or IRES sites, though expression of the dominant marker would then be limited to the germline (37–40). A second is that cargo genes incorporated into HEGs may be lost at appreciable frequencies. Tests of a replacement HEG carrying a cargo gene in Anopheles stephensi did not report such events (15). However, it is possible (species difference notwithstanding) that this again reflects the fact that these authors used a single gRNA in which cleavage results in two ends that are identical to the homology arms flanking the HEG. Perhaps this allows for more efficient complete homing, as compared with our multiplex gRNA scenario in which the ends generated by cleavage may need to be extensively chewed back in order to initiate HR with the HEG-bearing chromosome.

In our sequencing analysis of cleavage events, we also found that 5 out of 20 of analyzed flies had lost the dominant marker, but retained some fraction of Cas9 (functionality not tested) and another 5 flies carried only remnants of the gRNA cassettes but no Cas9 nor *td-tomato* (Fig. 3). Thus, not only are the observed homing rates too low for a HEG to bring about population suppression (3, 5, 11), but the HDR process, at least in our HEG configuration in *Drosophila*, is error prone. Incomplete homing is not fatal in the case of a suppression HEG, since it still creates a non functional copy of the gene. However, it can prevent effective population replacement if the HEG is meant to carry a cargo gene into the populations (15). Finally, we also observed recombination events involving the gRNA cassettes in 5 of 20 analyzed events, showing that the repetitive sequences in our multi promoter gRNA design are also prone to undergo recombination, with sequences in between them getting lost in the process. The repeats can be reduced by using polycistronic gRNAs expressed from a single promoter (41), though this may result different expression (processing) levels for the different gRNAs (42). In addition, the gRNA scaffolds would still have some level of repetitiveness. Thus, it remains to be shown if this approach will result in more stable copying of the HEG during the HDR process.

How can the frequency of complete homing events be increased? First, the NHEJ pathway can be suppressed (43). Second, Cas9 activity can be limited to stages of the cell cycle during which HDR is the dominant repair pathway by fusing cell cycle specific degrons to Cas9 (44, 45). It is possible that low rate of homing in *Drosophila* are (for unknown reasons) a species-specific problem, and that rates in important target species such as mosquitoes are higher (7, 12, 13, 15, 16). However, if this latter possibility is correct, it must have a genetic basis, which will be important to understand since polymorphisms in the relevant genes would provide a basis for selection of drive suppressors. Finally, as discussed above, the relationship between the sequences at the cleaved ends and those present in the homology arms of the HEG need to be explored so as to maximize cleavage while still allowing for a high frequency of complete homing. It will be particularly interesting to explore the consequences of locating multiple target sites within a small region.

### Spreading of the HEG through the female germ line is severely limited

An important problem encountered in our work is maternal deposition and activity of HEG components, i.e. Cas9 and gRNAs. Similar problems were observed and discussed previously, but with a focus on resistant allele generation (9, 11, 15–17). However, in the case of a suppression HEG germline carryover is devastating since most offspring coming from HEG-bearing females, including HEG-bearing heterozygotes, show loss-of-function phenotypes in somatic tissues. Thus, when the *Dfd*-HEG came from a female in the cross *Dfd*-HEG/+ x Df/TM3, Sb, the number of progeny carrying what should be a wildtype balancer chromosome (clv/TM3, *Sb* and HEG/TM3, *Sb*) was only 8% of that obtained when the sexes were switched.. For the *yg*-HEG only 17% of flies that inherited the HEG from the mother, were fertile, again showing the negative effect on fitness due to maternal carry over. *Nanos* plays roles in somatic tissues in the nervous system of *Drosophila* (*46*). However, it is unlikely that somatic expression in the zygote contributes to the *yellow-g* sterile phenotype, since no fitness cost was observed in HEG/TM3, *Sb* progeny when the HEG came from a father. In addition, while nanos is expressed in the nervous system, there are no reports of its expression in somatic follicle cells. The high maternal carryover fitness costs were reflected in the results of our drive experiments, in which both HEGs decreased in frequency or were lost entirely despite significant levels of homing. To mitigate the problem of maternal carryover different promoters are needed. Expression of the HEG could be limited to the male germline since based on our observations and those of others (32), carryover of Cas9 from sperm into the zygote is minimal. However, modeling suggests that, for a female fertility or viability locus, the genetic load introduced into the population as a result of homing in males only is lower than with homing in the female or in both sexes (3). Thus, conditions that allow for adult germline-specific Cas9 activity in females or both sexes are likely to be more useful for suppression HEGs. It may be possible to restrict Cas9 activity to oogenesis through the use of promoters that drive expression only during early stages of oogenesis, though HR-dependent repair from the homologous chromosome needs to be activity during these stages. Degrons could also be incorporated into the Cas9 protein to give it a shorter half life or modulate the phase of the cell cycle in which it is expressed, though Cas9 levels must not be reduced to such an extent that cleavage is no longer efficient. Cas9 inhibitor proteins could potentially also be used to temporally limit Cas9 activity, if they were expressed under the control of a late oogenesis-specific promoter, so as to inhibit Cas9 activity during early embryogenesis, after cleavage and homing had already occurred during oogenesis (47, 48).

To summarize, our results show that while gRNA multiplexing can prevent the appearance of alleles resistant to homing, a number of other issues remain to be addressed in order for HEG-based population suppression to become a reality. These include which germline promoters and other strategies to use to limit carryover and at the same time promote HR; how to increase the frequency of HR-dependent repair versus NHEJ; how to guarantee equal loading of members of a multiplex set of gRNAs; how to locate target sites and gRNAs so as to maximize the probability of multiple cleavage events, while also maximizing the probability of HR that results in complete homing events. In approaching these questions, particularly in the context of suppression HEGs, loci whose activity is required in adults for female fertility, such as *yellow-g*, are particularly useful. First, females lacking *yellow-g* are viable, but sterile with an easily scored collapsed egg phenotype. Second, because chromosomal deficiencies that lack *yellow-g* are available, females can be isolated that only carry sequences from the cleaved/homed allele, making it straightforward to characterize sequence changes at the locus. Third, *yellow-g* is conserved in other diptera including *Drosophila* pests such as *Drosophila* suzukii, and disease vectors such as mosquitoes (16). Thus, understanding how to effectively home at high frequency into *yellow-g* in one species may provides insights that can be translated to other species.

## Methods

### Target gene selection and gRNA design

We chose two target genes for the suppression HEG drives, the HOX gene *Dfd* (27) and *yellow-g* (26). The mosquito homolog of *yellow-g* has been tested for homing and suppression drive using a single gRNA previously (16). Disruption of *Dfd* results in embryonic lethality, while disruption of *yellow-g* results in defects in egg shell formation, resulting in female sterility. is involved in egg shell formation, thus, resulting in female-specific sterility. Potential gRNAs were ranked by on-target activity (49) in the benchling software suite (www.benchling.com) and selected to be spaced out over target gene exons in conserved regions (see Fig. 1A)

### Cloning of HEG constructs

#### guideRNAs

The starting construct pCFD3-dU6:3gRNA was a gift from Simon Bullock (Addgene plasmid # 49410) (29). The *BbsI* cut sites to insert a gRNA into the scaffold were replaced with *BsmBI* sites by digesting with *BbsI* and ligating annealed oligos as described on http://flycrispr.molbio.wisc.edu (FWD oligo: GTCGGGAGACGGACGTCTCT, REV oligo: AAACAGAGACGTCCGTCTCC). The *vermilion* marker gene was replaced by digesting with *HindIII* and ligating in a *white* marker gene.

We started by replacing the sgRNA scaffold with an optimized one published previously (30) in which the T base at position 4 was mutated to a G and the duplex was extended by 5 bp. The plasmid described above was digested with XbaI and BglII. Two fragments were amplified with the same plasmid as the template using primers 1-opti-g-FWD1 + 2-opti-g-REV1 and 3-opti-g-FWD2 + 4-opti-g-REV2. The nucleotide changes mentioned above were introduced in the primer overhangs of 2-opti-g-REV1 and 3-opti-g-FWD2. The construct was assembled in a two fragment Gibson assembly (50) to yield plasmid pCFD3-dU6:3gRNA-BsmBI-white-optimized-sg-scaffold.

Next, we added a second sgRNA promoter similar to pCFD4-U6:1_U6:3 tandem gRNAs (29). The pCFD3-dU6:3gRNA-BsmBI-white-optimized-sg-scaffold was digested with *XbaI*. The U6:1 promoter was amplified from genomic DNA extracted from a *w*-stock using the Qiagen DNeasy Blood & Tissue Kit with primers 5-U6:1-FWD1 + 6-U6:1-REV1. The sgRNA scaffold was amplified from the uncut plasmid using primers 7-U6:1-FWD2 + 8-U6:1-REV2. The final plasmid was assembled in a two fragment Gibson assembly yielding pU6:3-U6:1-tandem.

pU6:3-U6:1-tandem was used to subclone all guide RNAs into the sg-scaffold using the cloning strategy described by Port et al. (29). The pU6:3-U6:1-tandem was digested with *BsmBI* and the two guideRNAs were inserted in a one fragment Gibson reaction with the gRNA sequences encoded in the primer overhangs.

For each of the HEG constructs two of these guide cassettes were assembled for a total of four guides per construct:

p-yelG-left with primers 9-yelG-guide1-left FWD + 10-yelG-guide2-left REV, p-yelG-right with primers 11-yelG-guide4-right FWD + 12-yelG-guide3-right REV, p-dfd-left with primers 13-dfd-guide-left-fwd + 14-dfd-guide-left-rev and p-dfd-right with primers 15-dfd-guide-right-fwd + 16-dfd-guide-right-rev.

#### HEG construct

The HEG constructs utilized the *nanos* promoter and 3’ UTR to drive expression of Cas9. pnos-Cas9-nos was a gift from Simon Bullock (Addgene plasmid # 62208) (29). The mini-white marker in pnos-Cas9-nos was excised by digesting with ApaI and replaced with a *3xp3-td-tomato* marker cassette (51, 52). he nos-Cas9-nos cassette was then flanked by gypsy insulators by first cutting with NheI and inserting a gypsy insulator on the left and second, by cutting with SacII and inserting the insulator on the right. Finally, the whole insulated nos-Cas9-nos cassette was excised with *SapI* and *NsiI* and inverted in a one fragment Gibson reaction.

The resulting plasmid was digested with SnaBI and the left homology arm along with the left U6-tandem cassette was inserted in a two fragment Gibson reaction. Following that, it was digested with *ApaI* and the right homology arm and the right U6-tandem cassette were inserted as above (see Fig. S1 for a detailed cloning scheme and primers used).

#### HEG constructs without Cas9

To generate HEG fly strains that could be easily modified, we created versions of our HEG constructs in which the Cas9 cassette was replaced by an attP landing site. Flies carrying these constructs were also used as control strains in the drive experiments.

The two HEG constructs, *Dfd*-HEG and *yg*-HEG, were digested with HindIII to remove nos-Cas9-nos and the *3xp3-td-tomato* marker. The new construct was assembled in a three fragment Gibson reaction in which *td-tomato-sv40* expression was driven by the flightin promoter, and an attP landing site was included, resulting in the inactive HEG constructs, inactive-*Dfd*-HEG and inactive-*yg*-HEG.

### Fly germline transformation

The two HEG constructs *Dfd*-HEG and *yg-*HEG, as well as the two inactive HEG constructs, split-*Dfd*-HEG and split-*yg*-HEG, were injected into *w*-flies along with pnos-cas9-nos (29) as an additional source of Cas9. All injections were carried out by Rainbow Transgenic Flies. The injected G_0_ flies were outcrossed to *w*- and the resulting progeny were screened for the fluorescent *3xP3-td-tomato* eye marker. Transformed flies were kept as heterozygous stocks by outcrossing some male flies to *w*-each generation.

### Fly crosses

To analyze homing and cleavage rates of the generated HEG strains we set up the following crosses with deficiency lines of the targeted gene. 25 Heterozygous HEG bearing females were crossed individually to deficiency/Balancer males and vice versa. The *Dfd*-HEG strain was crossed to Df(3R)*Dfd*13/TM3, *Sb* (Bloomington #1980) (27):

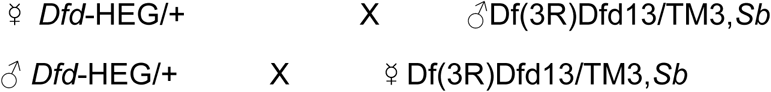

The resulting progeny of the single fly crosses were scored for possible genotypes: *Dfd*-HEG/TM3, *Sb*; +(or clv)/TM3, *Sb*; +(or clv)/Df(3R)*Dfd*13 and Df(3R)*Dfd*13/*Dfd*-HEG.

The *yg*-HEG strain was crossed to Df(3L)BSC384/TM3, *Sb* (Bloomington #24408) (53):

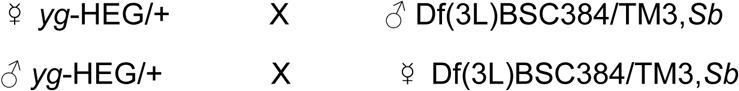

The resulting progeny was scored for possible genotypes: *yg*-HEG/Df(3L)BSC384; *yg*-HEG/TM3, *Sb*; + (or clv)/ Df(3L)BSC384 and +(or clv)/TM3, *Sb*. The flies were also sexed in this case, because only females elicit the *yellow-g* phenotype. All these females were outcrossed to *w*-males subsequently to check for sterility.

Females coming from a HEG bearing father with the genotypes +(or clv)/TM3, *Sb* and yg-HEG/TM3, *Sb* were also outcrossed to check for paternal carry over of the HEG components. Only five vials were scored, showing no obvious effects on fertility.

### Homing- and cleavage rate calculations

For both drives the homing rate can be estimated by looking at the proportion of HEG/TM3, *Sb* individuals out of all TM3, *Sb* individuals since no TM3, *Sb* individuals should (in the absence of carryover) have died as a result of HEG activity. As such, the average or expected number of individuals per genotype can be estimated as half of the sum of the number of the HEG/TM3, *Sb* and +/TM3, *Sb* individuals, a quantity we shall denote as G. If we assume an equal number of offspring for each genotype, we get our estimated homing rate by taking the HEG/TM3, *Sb* individuals, subtracting G, and then dividing the difference G. The way to intuit this is as follows: HEG/TM3, *Sb* is a sum of the individuals that inherited the HEG through Mendelian transmission and those who received it from a homing event. Thus by subtracting G and dividing the difference by G, we estimate the percentage of the HEG/TM3, *Sb* individuals that received the HEG from a homing event rather than through normal inheritance (see homing rate equation below).

Similarly, since the only way to obtaint +/Df viable individuals (or fertile females in the case of the yG-HEG) is if cleavage does not occur t, we can use the frequency of these events to estimate the cleavage rate for both drives by subtracting +/Df from G and dividing the difference by G (see cleavage rate equation below).

Note that for the yG-HEG the +/TM3, *Sb* pool represents both +/TM3, *Sb* and clv/TM3, *Sb* since we can’t distinguish between the two phenotypically.

*G* = ((*HEG*/*TM* 3, *Sb*) + (+ /*TM* 3, *Sb*))/2

*homing rate* = (*HEG*/*TM* 3, *Sb* − *G*) / *G*

*cleavage rate* = (*G* − (+ /*Df*)) / *G*

### Sequence analysis of escapers

We sequenced the targeted region from progeny flies that survived (*Dfd)* or were fertile (*yellow-g*) and that carried the Df chromosome, from parental crosses that were HEG/+ x Df/TM3, *Sb*. These were identified as being Df-bearing by virtue of the fact that they lacked the balancer chromosome TM3, *Sb.* Genomic DNA was extracted from single flies using the Qiagen BloodNEasy kit. The genomic regions containing guide RNA target sites were amplified using NEB Longamp PCR mastermix. Because of the long intron, the *Dfd* genomic region was amplified in two fragments. Fragment 1 containing gRNA 1 and 2 was amplified with primer 47-dfd-left-fwd and 48-dfd-left-rev. Fragment 2 containing gRNA 3 and 4 was amplified with primer 49-dfd-right-fwd and 50-dfd-right-rev. The same primers were used for sequencing. The *yellow-g* genomic region was amplified with primers 51-yG-fwd and 52-yG-rev. These primers were also used for sequencing along with an additional primer 53-yG-mid.

### Sequencing of cleavage events

We selected ten sterile female progeny of the genotype clv/Df from crosses of female HEG/+ and male HEG/+ flies to individuals that were Df/TM3, *Sb*. Genomic DNA from single flies was extracted using the Qiagen BloodNeasy kit. The targeted genomic region was amplified with NEB Longamp PCR Mastermix using primers located outside of the homology arms used for the homing construct (54-yG-outside FWD + 55-yG-outside REV). The resulting amplicons were purified on a gel and sequenced with primers 56-yG-seq1, 57-yG-seq2, 58-yG-seq3 and 59-yG-seq4.

### Drive experiments

Drive experiments were set up in fly food bottles for the two active homing stocks *Dfd*-HEG and *yg*-HEG, as well as for the two inactive versions inactive-*Dfd*-HEG and inactive*-yg*-HEG as the controls. To seed the bottles we crossed 30 female *w*-virgins to 30 heterozygous males of the corresponding drive or control stocks. After allowing the flies to mate for two days they were transferred to a fresh fly food bottle together with 30 *w*-females that had been similarly mated with *w*-males. All drive experiments were set up in four biological replicates.

The flies were allowed to lay eggs for two days in the fresh bottle before being removed. After the next generation of adult flies hatched and mated in this bottle, all flies were dumped on a CO_2_ pad and scored for the marker of the homing construct under a fluorescent microscope. For the bottles to not become overcrowded we set an artificial population size limit of 200 flies. These were transferred to a new bottle as the next generation and the cycle was repeated for 12 generations.

### Modeling

We use a deterministic, discrete-generation population frequency framework to model the spread of each HEG through a population, assuming random mating. This model is a variant of one used previously (54). It consists of a series of difference equations used to calculate the expected frequency of each genotype based on the frequencies of all genotypes from the previous generation, augmented by fitness effects, cleavage and homing events, and maternal carryover, and is finally adjusted by a normalization factor. These equations, while straightforward, are rather lengthy due to the number of possible crosses, so we have provided them directly within our model in a supplementary matlab file (File S1).

### Containment of HEG flies

All HEG bearing flies were contained following recommended barrier confinement strategies (55). Flies were kept in triple nested containers and kept in a dedicated room behind 3 Doors. Doors were additionally fitted with nets and fly traps were placed in the room. One investigator (G.O.) performed all experiments and fly handling. All flies were were autoclaved prior to disposal.

### Figures

Fig. 2 and Fig. 5 were plotted in R using the “ggplot2” and “RColorBrewer” packages (56–58). All figures were assembled and labeled in Inkscape.

## Acknowledgements

We thank Marlene Biller for technical assistance. This work was supported by a grant from the California Cherry Board. G.O. was supported by a research fellowship from the German Research Foundation GO 428/1-1. T.I. was supported by NIH training grant 5T32GM007616-39.

## Author contributions

G.O. and B.A.H designed research; G.O. performed research; G.O., T.I., and B.A.H. analyzed data; T.I. modeled the drive, G.O., T.I., and B.A.H. wrote the paper.

## Supporting Information

**Fig S1: Cloning strategy**

**Table S1: Genotype counts of crosses to the deficiency line**

**Table S2: Drive experiment counts**

**Table S3: List of primers**

**File S1: Matlab source file of model**

**Fig S1:**
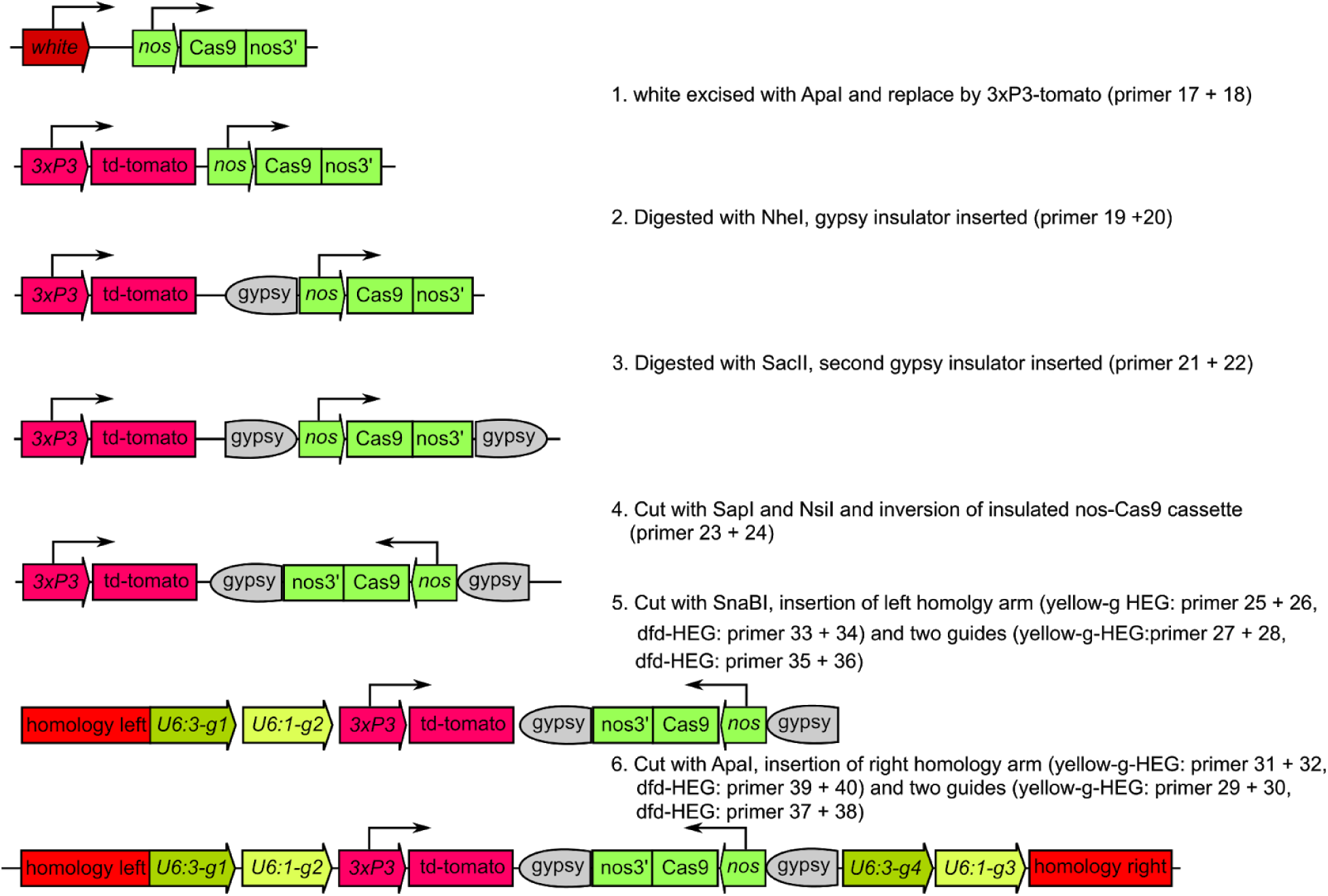
Cloning strategy for HEG and split-HEG constructs. Step by step assembly guide listing used enzymes and primers. See Material and Methods section for details.

